# Large temperature excursions have modest impacts on community composition in the high diversity gut microbiome of omnivorous American cockroaches (*Periplaneta americana*)

**DOI:** 10.64898/2026.01.21.700893

**Authors:** Kevin C. Riedmuller, Josey E. Dyer, Elizabeth A. Ottesen

## Abstract

Microbial residents of ectothermic hosts are exposed to variations in temperature that have the potential to impact their physiology and the host-microbe symbiotic relationship. In this experimental warming study, laboratory populations of American cockroaches (*Periplaneta americana*) were kept at a baseline low room temperature of 20-22°C or a high temperature of 30°C for two weeks. We quantified bacterial load and performed high-throughput 16S rRNA gene sequencing to assess the hindgut microbiome’s response to a near 10°C shift in environmental temperature. We report modest impacts of temperature on cockroach gut microbiome composition. The high temperature treatment induced increases in the relative abundance of Proteobacteria and Euryarchaeota phyla as well as the Lactobacillaceae and Enterococcaceae families. We also observed increased interindividual variability. There were no significant differences in the dominant Bacteroidota or Firmicutes phyla and no significant losses or reductions in taxa or bacterial load, respectively. This suggests that the gut community of American cockroaches is largely resilient to prolonged increases in temperature and has implications for the cockroach to withstand the impacts of climate change.

**Importance:** Insects, as with most animals, often harbor microbial symbionts that play an essential role in host health and nutrition. As insects are ectotherms, these microbial symbionts are subject to the same temperature fluctuations as their hosts, potentially impacting host temperature responses. Here, we demonstrate that the American cockroach (*Periplaneta americana*) gut microbiome exhibits only modest changes following an ∼10°C increase in environmental temperature. This contrasts with studies in other insects, whose microbiota were highly responsive to temperature variation. This work illustrates that the microbiota of insects may vary in their sensitivity to long-term temperature shifts, providing a more comprehensive understanding of potential variability in insect responses to climate change.

## Introduction

Insects are some of the most abundant multicellular organisms on earth with an estimated 5.5 million species (1). They are key drivers of ecological and biological processes ranging from nutrient cycling (2,3) to disease dynamics (4,5) and agricultural productivity (6,7). As ectotherms, environmental temperature strongly governs their biological timing, physiology, distribution, and fitness (8). Consequently, changes in environmental temperature can have outsized and varied impacts on insects’ roles in these critical systems. For example, simulated warming in mesocosms containing *Formica subsericea*, an ant native to eastern North America, led to decreased decomposition rates and nitrogen availability (9). In contrast, a field study of *Formica Manchu* nests on the southeastern Tibetan Plateau in China found that decomposition rates and nitrogen availability remained stable under simulated warming conditions (10). As 2024 was the warmest year on record and global surface temperatures are rising faster than any other period over the last 2000 years (11,12), understanding how insects are affected by warming climate has become increasingly important.

Insects have evolved a wide variety of behavioral and physiological thermoregulatory responses for survival that modulate metabolism, stress response, and growth and development (8). Studies frequently report faster growth and/or development rates in response to increased temperature, which is often accompanied by a decrease in body size (13–15). However, one aspect of insect adaptation to temperature that is less well-understood is the impact of temperature on the insect gut microbiome. The gut microbiome plays a critical role in host nutrition (16–18), development (19–21), and immunity (22–24). As symbionts of ectotherms, gut microbes (and the services they provide to their hosts) are subject to changes in environmental temperature alongside their hosts. Studies in three species of stink bugs have reported reduction or complete loss of gut bacterial symbionts in response to increased environmental temperature by as little as 2.5°C (25,26). Similarly, a paired observational and experimental study in desert arboreal ants found that increased environmental temperature resulted in decreased bacterial loads as well as moderate shifts in microbial community composition. Notably, these changes in the gut microbiome were due to only a 1°C shift from baseline temperature in the experimental treatment (27). These data parallel studies in other ectotherms like corals, sponges, and mussels where increased temperature saw substantial shifts, reductions, or complete loss of symbiotic microbes (28–30). Overall, studies in a variety of invertebrates thus far tend to observe marked changes in both high and low diversity gut microbiomes and subsequent decreases in host fitness as a result of elevated temperatures ranging from ∼1-9°C from baseline temperatures.

Omnivorous cockroaches carry a high-diversity hindgut microbiome (31–33) which is believed to play an important role in facilitating host digestion of complex polysaccharides and other refractory dietary components (34,35). The cockroach gut microbiome has been shown to play a vital role in shaping host development (36,37), immune activity (38,39), and behavior (40). It also shows evidence of long-term co-evolution with its host (41) and likely plays a key role in their effectiveness as decomposers and scavengers—functions that contribute significantly to global nutrient cycling due to their ubiquity across terrestrial ecosystems. For example, in central Amazonian inundation forests, at least eight different species of cockroaches were identified, one of which was estimated to consume up to 6% of annual leaf litter (42). Their effectiveness as decomposers and scavengers also contribute to pest species’ ability to infest virtually all human spaces (43). Pest species of cockroaches pose significant human health risks as they are a common cause of allergy and asthma (44,45) and vectors for a variety of pathogens (46,47). Moreover, they contribute to economic losses by driving up medical costs (48), destroying property and goods (43), and necessitating continual pest control (49). Given their ecological and economic importance, it is critical that we understand the effects of warming temperatures on the cockroach and its closely associated gut microbiome.

While little is known about the impact of temperature on the cockroach gut microbiome, studies of their close relatives the termites (50) show marked temperature impacts. An experimental study in *Reticulitermes flavipes* comparing the gut microbiota at 15°C, 27°C (baseline) and 35°C found dramatic shifts in gut microbiome composition. Colder temperatures were associated with increased relative abundances of Proteobacteria while heat stress was associated with decreased bacterial richness, loss of protist associated bacteria and increased relative abundances of Bacteroidota (51). In contrast, a field study in Janesville, Wisconsin between July and December observed only modest seasonal shifts in gut community composition of *R. flavipes*. Notably, the mean environmental air temperature during the collection period ranged from 22°C to −2°C, which was substantially lower than the warming treatment in the experimental study (52).

The aim of this work was to investigate the impacts of environmental temperature on the gut microbiome of the model insect *Periplaneta americana*. It has been observed that their thermal preference ranges from ∼24-33°C (53). However, in many cases experimental insects are kept at a variety of temperatures within or below the thermal preference range (54–56) or at an unspecified room temperature (33,57–59). While such experimental differences likely make little difference to the gut microbiota of endotherms such as mice, it poses a potential complicating factor in studies of microbial symbioses of ectotherms. As a result, we sought to examine the impact of temperature on the gut microbiota of omnivorous cockroaches. To do so, we employed bacterial load quantification and high throughput 16S rRNA gene sequencing to assess the hindgut microbiome’s response to an increase in environmental temperature from a ‘low’ laboratory room temperature of 20-22°C to 30°C, a ‘high’ temperature used to speed reproduction and development (60). This work provides insight into how temperature can impact the gut microbiota of ectothermic hosts; This knowledge has implications for both organismal responses to climate shifts and experimental practice for laboratory investigation of ectotherm symbioses.

## Materials and methods

### Insects and experimental design

*P. americana* cockroach stock colonies have been maintained in captivity at room temperature (20-22°C) for over 15 years. Stock colonies live in 10-gallon glass aquarium tanks with wood chip bedding (PJ Murphy Forest Products SaniChips) and cardboard tubes for housing. They were provided dog chow (Purina One) and water (via cellulose sponge) *ad libitum*.

For temperature treatments, 20-24 mixed-sex, adult cockroaches were selected from stock colonies, divided into two plastic terrariums, and maintained under stock colony conditions at room temperature (20-22°C) or 30°C for 14 days. Dog chow with any visible mold growth was removed daily. The experiment was performed in duplicate. Cohort 1 and 2 treatments were completed in April of 2022; cohort 2 treatments were initiated one week after cohort 1 began. We also include data from a pilot experiment conducted November of 2020 (cohort 3). The room temperature treatment for this cohort was slightly warmer, ranging from 22-25°C.

### Sample collection and DNA extraction

Cockroaches were placed on ice in sterile petri dishes until torpid (∼5 min). The entire gut (foregut to rectum) was excised with a scalpel and forceps; any visible fat body and exoskeleton was removed. The gut tract was then moved to a sterile aluminum dish, placed over dry ice, and the hindgut was resected. Hindguts were transferred to 500 µL of sterile 1x phosphate buffered saline and stored at −20°C.

Hindguts were thawed and homogenized with a sterile microcentrifuge pestle. DNA was extracted from a 200 µL aliquot of sample homogenate using the E.Z.N.A Bacterial DNA Kit (Omega Bio-tek, Norcross, GA) per manufacturer’s protocol with the following modifications: 30 min lysozyme incubation, optional glass bead beating step included, and RNase A step excluded. Sample DNA was eluted in 50 µL of elution buffer. DNA concentrations and A260/280 values were measured with a BioTek Synergy HTX Multimode plate reader (Agilent, Santa Clara, CA). Sample DNA was stored at −20°C and concentrations ranged from 30 to 300 ng/µL (Table S1).

### Bacterial load quantification

Total number of bacteria per sample was assessed via qPCR of the 16S rRNA gene as described in Nadkarni et al. (61). qPCR was performed in triplicate, 10µL reactions using a QuantStudio 6 flex instrument and QuantStudio Real Time PCR Software (v1.3) (Applied Biosystems, Waltham, MA). Reactions consisted of 1x TaqMan Multiplex Master Mix (Applied Biosystems), 0.3 µM forward primer 5‘-TCCTACGGGAGGCAGCAGT-3’ (Tm, 59.4 °C), 0.3 µM reverse primer 5 ‘-GGACTACCAGGGTATCTAATCCTGTT-3’ (Tm, 58.1 °C), 0.25 µM hydrolysis probe (6-FAM)-5 ‘-CGTATTACCGCGGCTGCTGGCAC- 3 ‘-(TAMRA) (Tm, 69.9 °C), and 1µL sample DNA (see Table S1 for DNA quantities) under the following conditions: 95°C for 20 s followed by 40 cycles of 95°C for 1 s and 60°C for 60 s. A standard curve was generated with a 10-fold dilution series of gBlocks Gene Fragment of the *Frigididesulfovibrio cuneatus* 16S gene (GenBank: X99503.1, position 330-798) with a linear dynamic range of 10-10^8^ copies. The slope, y-intercept, and r^2^ of the standard curve was −3.529, 41.67, and 0.994, respectively. Amplification efficiency was 92.04%.

qPCR of a *Blattabacterium* MIP/aquaporin family protein coding sequence was performed to account for the cockroach endosymbiont’s contribution to bacterial load. Triplicate, 10 µL reactions consisted of 1x TaqMan Environmental Master Mix 2.0 (Applied Biosystems), 0.1 µM forward primer 5 ‘-CAGCTAATGCCCATCCCATAG-3’ (Tm, 55.4 °C), 0.1 µM reverse primer 5 ‘-GGAAATGGAGTGGTGGCTAAT-3’ (Tm, 55.0 °C), 0.1 µM hydrolysis probe (6-FAM)-5 ‘-TTGTGACCCATCCACCATCTCCAT-3 ‘-(MGB-NFQ) (Tm, 63.7 °C), and 1µL sample DNA (see Table S1 for DNA quantities) under the following conditions: 95°C for 10 min followed by 40 cycles of 95°C for 15 s and 60°C for 60 s. A standard curve was generated with a 10-fold dilution series of gBlocks Gene Fragment (CAGCTAATGCCCATCCCATAGTGATTGTGACCCATCCACCATCTCCATGTCCTTTTGTTTTAGACAATAGAACATTAGCCACCACTCCATTTCC) with a linear dynamic range of 10-10^8^copies. The slope, y-intercept, and r^2^ of the standard curve was −3.526, 38.97, and 0.993, respectively. Amplification efficiency was 92.15%.

C_q_ values for both assays were auto selected by the QuantStudio Real Time PCR Software (v1.3). All primers and gBlocks were synthesized by Integrated DNA Technologies with standard desalting purification. Hydrolysis probes were synthesized by Applied Biosystems. Primer set specificity was validated via endpoint PCR and agarose gel electrophoresis.

Hindgut bacterial load *L* was calculated as follows:

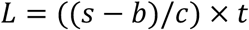

where *s* = 16S copy number, *b* = *Blattabacterium* MIP/aquaporin family protein copy number, *c* = DNA quantity in qPCR reaction (ng), and *t* = total sample DNA (ng). Reported 16S and *Blattabacterium* MIP/aquaporin family protein copy numbers are averages from triplicate qPCR reactions. Bacterial load boxplots were generated with the Tidyverse package (62).

### Library preparation and sequencing

The V4 region of the 16S rRNA gene from each hindgut sample was amplified using a two-step PCR protocol as described in Tinker et al. (63). Initial amplification of the V4 region was performed with a 10 µL PCR reaction that consisted of 1x Q5 hot start reaction buffer, 0.02 U/µL Q5 hot start high-fidelity DNA polymerase (New England BioLabs, Ipswitch, MA), 0.5 µM 515F primer 5 ‘-GTGCCAGCMGCCGCGGTAA-3 ‘, 0.5 µM 806R primer 5 ‘-GGACTACHVGGGTWTCTAAT-3 ‘, 200 µM deoxynucleotide triphosphates (dNTPs), and 0.8 ng/µL sample DNA under the following conditions: 98°C for 30 s, followed by 15 cycles of 98°C for 10 s, 52°C for 30 s, and 72°C for 30 s, followed by a final extension step at 72°C for 2 min. The product of reaction 1 was reamplified in a subsequent 30 µL PCR reaction, containing 1x Q5 hot start reaction buffer, 0.02 U/µL Q5 hot start high-fidelity DNA polymerase, 0.5 µM 515F barcoded primer, 0.5 µM 806R barcoded primer, 200 µM dNTPs, and 9 µL of reaction 1 product under the following conditions: 98°C for 30 s, followed by 4 cycles of 98°C for 10 s, 52°C for 30 s, and 72°C for 30 s, followed by 6 cycles of 98°C for 10 s and 72°C for 1 min, followed by a final extension step at 72°C for 2 min. All primers were synthesized by Integrated DNA Technologies with standard desalting purification (Coralville, IA).

PCR products were verified by agarose gel electrophoresis and purified with the E.Z.N.A Cycle Pure Kit (Omega Bio-tek) per manufacturer’s protocol and eluted in 30 µL elution buffer. Purified amplicon concentrations and A260/280 values were measured with a BioTek Synergy HTX Multimode plate reader. Amplicons were normalized and pooled to a final concentration of 10 ng/µL. Quality of the prepared amplicon library was assessed with the Agilent 2100 Bioanalyzer DNA-HS assay (Agilent) before submission to the Georgia Genomics and Bioinformatics Core for paired-end 250bp sequencing on an Illumina MiSeq platform (Illumina Inc., San Diego, CA).

#### Sequence processing

Demultiplexed, paired-end fastq files were analyzed with the standard DADA2 (64) pipeline. Forward reads were truncated at 240 bp and reverse reads were truncated at 160 bp. Samples were pooled during the sample inference step to increase sensitivity to sequence variants present at low abundances in multiple samples. Taxonomy was assigned to amplicon sequence variants (ASVs) with the DADA2 package and the Silva reference database (v138.1) (65); reads corresponding to *Blattabacterium* (cockroach endosymbiont located in the fat body), chloroplasts, and mitochondria were removed. All downstream analyses of sequencing and bacterial load data were performed using R Statistical Software (v4.2.1) (66) and Geneious Prime 2025.0.3 (https://www.geneious.com).

#### Alpha and beta diversity

All libraries were resampled to a depth of 10,797 reads with the Vegan package (v2.6-4) (67) for alpha and beta diversity metrics. Alpha diversity was assessed by calculating the Shannon diversity index, richness (number of observed taxa), and Pielou’s evenness (Shannon diversity index/log(richness)) (67). Alpha diversity was visualized with violin plots via the Tidyverse packages (v2.0.0) (62). Beta diversity was assessed by calculating weighted and unweighted (incidence based, equivalent to Sorenson index) (68) Bray-Curtis dissimilarities (67). Beta diversity was visualized with nonmetric multidimensional scaling ordinations and violin plots via the Tidyverse and Vegan packages (62,67).

#### Differential abundance analysis

DESeq2 (v1.38.3) (69) was implemented for ASV level differential abundance analysis to identify temperature responsive taxa. The ASV raw counts table was passed to the DESeq command with default parameters and ‘design = ∼ cohort + temperature’, testing for the effect of temperature while controlling for cohort variation. ASVs with an adjusted *p* value < 0.05 were considered significant (Table S2). A phylogenetic tree of differentially abundant ASVs present in the 12 most abundant families (maximum relative abundance > 10%) was generated in Geneious Prime. Sequences were aligned with Clustal Omega (v1.2.3) (70) and the phylogenetic tree was built using FastTree (v2.1.11) (71,72). The *Desulfovibrionaceae* clade was used as an outgroup for rooting, and tree visualization was performed in R with the Tidyverse and ggtree (v3.6.2) (73) packages. A heatmap of differentially abundant ASVs present in the 12 most abundant families was generated with the ComplexHeatmap package (74,75).

### Data availability

The sequences generated from this experiment were submitted to the NCBI Sequence Read Archive and are available under BioProject PRJNA1380472.

## Results

### Hindgut bacterial load is stable across temperature treatments

To assess the effect of temperature on hindgut bacterial load, we performed qPCR of the 16S rRNA gene with a universal primer/probe set as described in Nadkarni et al (61). To account for 16S rRNA copy contribution from the cockroaches’ *Blattabacterium* endosymbiont, we also performed qPCR of a *Blattabacterium* gene in the MIP/aquaporin family. Wilcoxon rank-sum tests, a non-parametric test comparing the distribution of independent groups, were used for two-group comparisons. Kruskal-Wallis tests, an extension of the Wilcoxon rank-sum test, were used for the overall comparison of three or more groups (non-parametric ANOVA analogue). Significant Kruskal-Wallis tests were followed up with post-hoc Dunn’s test with Bonferroni adjustment to determine which groups’ distributions were significantly different (FSA v0.9.6) (76). A Wilcoxon rank-sum test found no significant difference in hindgut bacterial load across temperature treatments (*p* = 0.26). A Kruskal-Wallis test found a significant difference in hindgut bacterial load among cohorts (*p* = 0.04). However, Dunn’s test did not indicate significant differences between cohorts: cohort 1 and 2 (*p* = 0.10), cohort 1 and 3 (*p* = 1.00), and cohort 2 and 3 (*p* = 0.07) (Fig 1).

**Fig 1.**
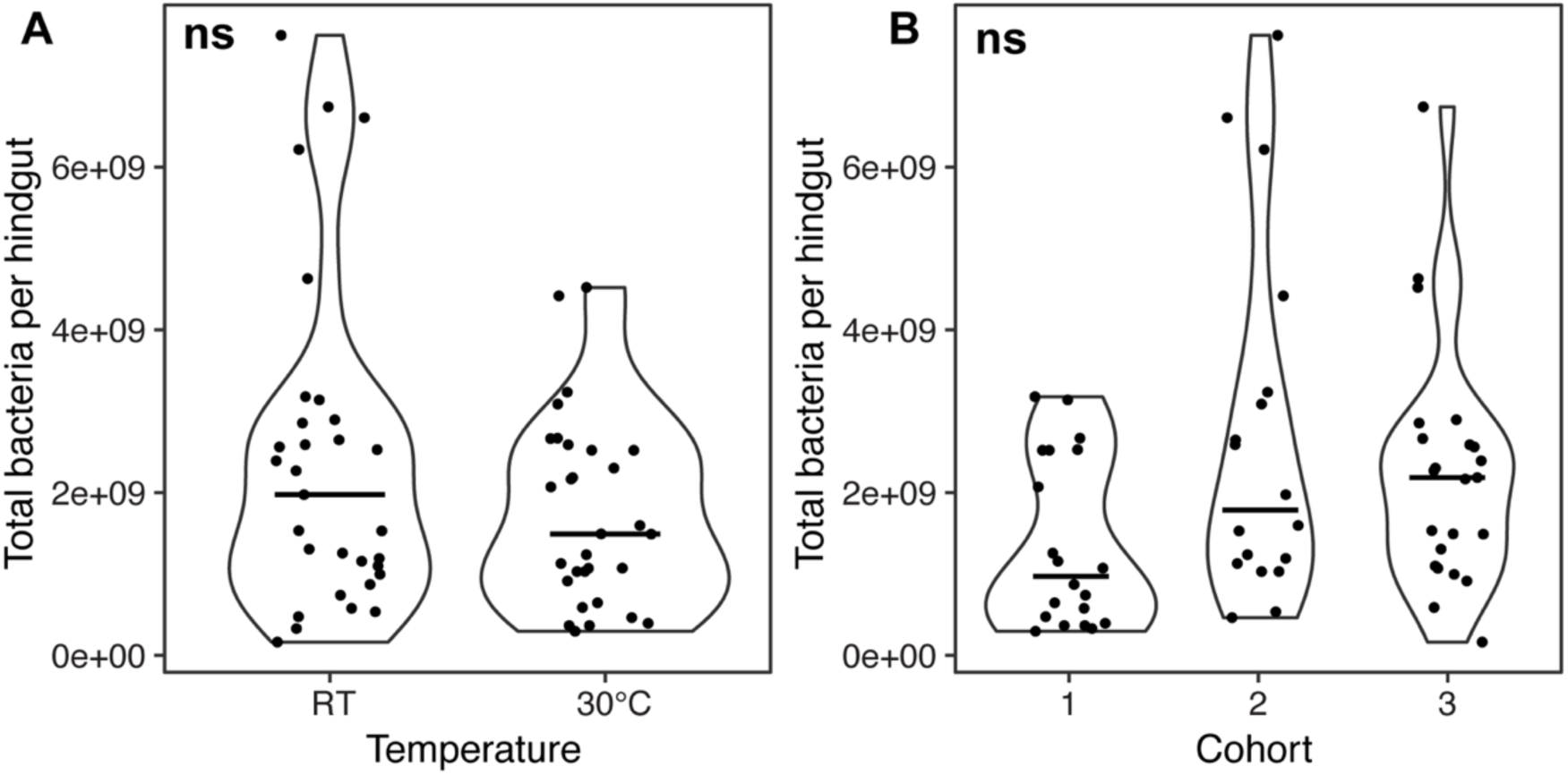
Comparison of hindgut bacterial load across temperature treatments and cohorts. Violin plots of hindgut bacterial load across (A) temperature and (B) cohort where bars represent the median and points represent individual samples. Wilcoxon rank-sum test was used to compare across temperature treatments. Kruskal-Wallis and post-hoc Dunn’s test with Bonferroni adjustment was used to compare across cohorts. *Blattabacterium* counts subtracted from total 16S rRNA copy number. Bacterial load normalized by total sample DNA. RT = room temperature, ns = no significance.

### Hindgut microbial community composition

A total of 4,526,160 rRNA gene sequences were obtained and 2,580,005 passed filtering parameters, resulting in an average of 42,295 reads per sample. One sample was removed prior to downstream analysis due low sequencing depth (Table S3). 2,923 unique ASVs were identified across all samples.

Overall gut microbiome composition exhibited modest changes in response to temperature. Wilcoxon rank sum tests found no significant differences in the Shannon diversity, Pielou’s evenness, or richness (total number of ASVs) of samples at the ASV or family level across temperature treatments (Shannon: ASV *p* = 0.86, family *p* = 0.57; Pielou’s: ASV *p* = 0.88, family *p* = 0.56; richness: ASV *p* = 0.64, family *p* = 0.47) (Figs 2A and S1A). However, beta diversity measures suggest modest but significant impacts of temperature on gut microbiome community composition. Dunn’s test indicated that between-temperature treatment dissimilarities were significantly greater than within- room temperature treatment dissimilarities at the ASV (*p* < 0.001) and family level (*p* < 0.001) (Figs 2B and S1B). While nonmetric multidimensional scaling (NMDS) of weighted Bray-Curtis dissimilarities did not indicate clear separation of samples by temperature, permutational multivariate analysis of variance (PERMANOVA) (67) found a significant effect at the ASV (*p* = 0.001, r^2^ = 0.05) and family (*p* = 0.001, r^2^ = 0.05) levels (Figs 2C and S1C). Temperature was also observed to have impacts on beta dispersion, with the 30°C treatment having higher within-group dissimilarity than the room temperature treatment (Dunn’s test: ASV *p* < 0.001, family *p* < 0.001) (Figs 2B and S1B).

**Fig 2.**
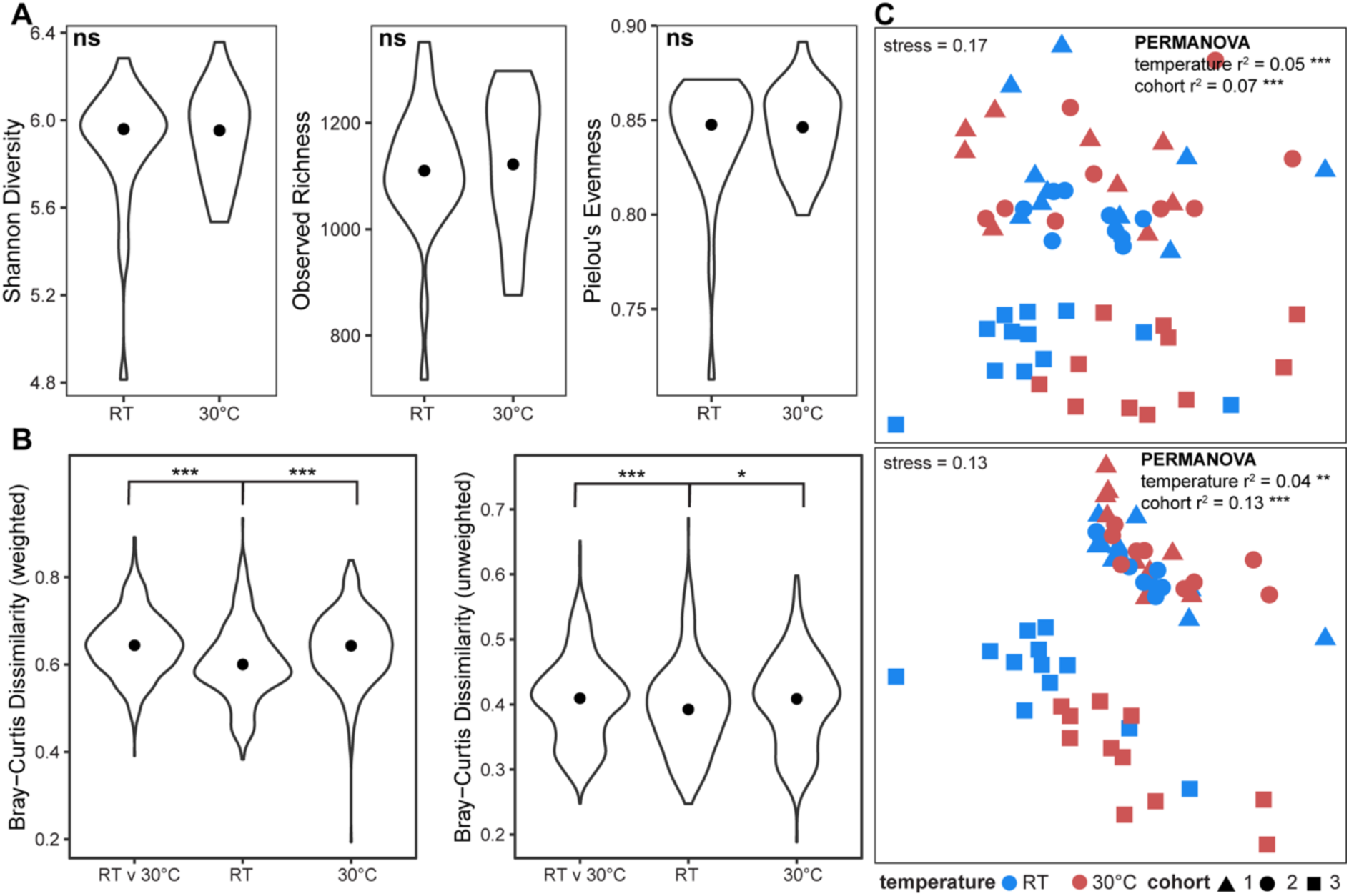
ASV level comparison of hindgut microbial community composition across temperature treatments. (A) Violin plots of alpha diversity measurements (Shannon diversity, observed richness, Pielou’s evenness) where points represent the medians. (B) Violin plots of weighted (left) and unweighted (right) Bray-Curtis dissimilarities within and between temperature treatments where points represent the medians. Wilcoxon rank-sum tests were used to compare alpha diversity measures. Kruskal-Wallis and post-hoc Dunn’s test with Bonferroni adjustment was used to compare Bray-Curtis dissimilarities. (C) Nonmetric multidimensional scaling (NMDS) of weighted (top) and unweighted (bottom) Bray-Curtis dissimilarities. NMDS stress was calculated with the metaMDS() function from the Vegan package. PERMANOVA was used to calculate r^2^ and *p* values for temperature and cohort. All libraries were resampled to a depth of 10,797 reads. RT = room temperature, * = *p* < 0.05, ** = *p* < 0.01, *** = *p* < 0.001, ns = no significance.

Given the large time gap between experiments, we also examined cohort impacts on community composition. Notably, cohort 3 samples tended to cluster separately from other cohorts by weighted and unweighted Bray-Curtis metrics and PERMANOVA found a significant effect for cohort on community composition at the ASV (weighted: *p* = 0.001, r^2^ = 0.07; unweighted: *p* = 0.001, r^2^ = 0.13) and family (weighted: *p* = 0.04, r^2^ = 0.04, unweighted: *p* = 0.001, r^2^ = 0.22) level (Figs 2C and S1C). Dunn’s test indicated significantly greater between-cohort dissimilarity for all comparisons involving cohort 3 compared to the between-cohort dissimilarity of 1 and 2 by weighted and unweighted metrics at the ASV level (*p* < 0.001) (Fig S2). This suggests that some observed taxa in cohort 3 differ from the observed taxa in cohorts 1 and 2. This trend aligns temporally with each experiment, where cohort 1 and 2 treatments were carried out within a week of each other while cohort 3 was carried out over one year earlier. This indicates minor, detectable drift in taxonomic composition of our stock colonies. To investigate whether cohort effects were obscuring temperature responses, we examined temperature effects on gut microbiome composition separately for cohorts 1 and 2, which were combined due to their more similar community composition, and cohort 3 (Figs 2C and 3). Accounting for cohort variation, samples separated more clearly by temperature at the ASV level (weighted Bray-Curtis dissimilarity) than at the family level (Fig 3), although the effect size of temperature calculated by PERMANOVA was still modest (cohort 1 and 2: *p* = 0.003, r^2^ = 0.07, cohort 3: *p* = 0.001, r^2^ = 0.13). This suggests that, while we observe stability of the gut microbiome at the family-level in response to temperature, there may be some ASV-level differences contributing to observed stability.

**Fig 3.**
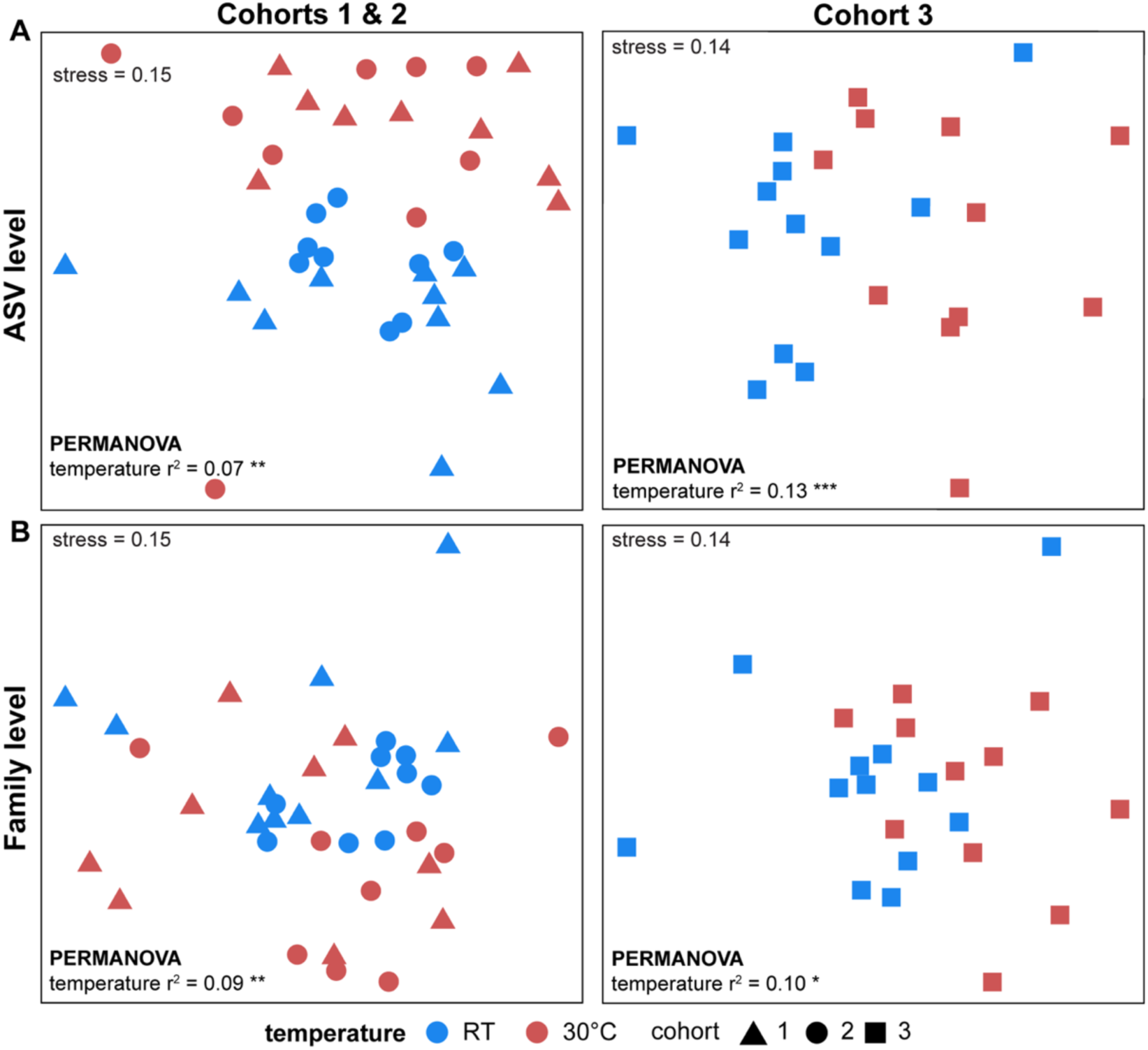
ASV and family level comparison of Bray-Curtis dissimilarities accounting for cohort differences. Nonmetric multidimensional scaling (NMDS) of weighted Bray-Curtis dissimilarities at the (A) ASV and (B) family level. NMDS stress was calculated with the metaMDS() function from the Vegan package. PERMANOVA was used to calculate r^2^ and *p* values for temperature and cohort. All libraries were resampled to a depth of 10,797 reads. RT = room temperature, * = *p* < 0.05, ** = *p* < 0.01, *** = *p* < 0.001.

### Taxa exhibiting significant responses to temperature differences

All ASVs present in more than 50% of the room temperature samples were also present in the 30°C samples, and vice versa, suggesting that temperature differences in community composition were primarily the result of changes in ASV relative abundance. We performed differential abundance analysis with DESeq2 to further explore ASV level responses to environmental temperature. Accounting for cohort variation, 231 ASVs were identified as significantly differentially abundant (*p* < 0.05) across temperature treatments (Table S2). Of those 231 ASVs, 123 belonged to the most abundant families (maximum relative abundance > 10%) in our samples (Figs 4 and S3). Ruminococcaceae and Lachnospiraceae families had a comparable number of ASVs enriched in both temperature treatments. This may be indicative of ASV-level switching among functionally redundant taxa within these families. However, we did not see this trend for the remaining families. All but one Bacteroidaceae ASV was enriched in the room temperature treatment and all Lactobacillaceae and Enterococcaceae ASVs were enriched in the 30°C treatment. We found that most differentially abundant ASVs in the Tannerellaceae, Dysgonomonadaceae, Rikenellaceae, Christensenellacae, and Desulfovibrionaceae families were enriched in the 30°C treatment (Figs 4 and S3).

**Fig 4.**
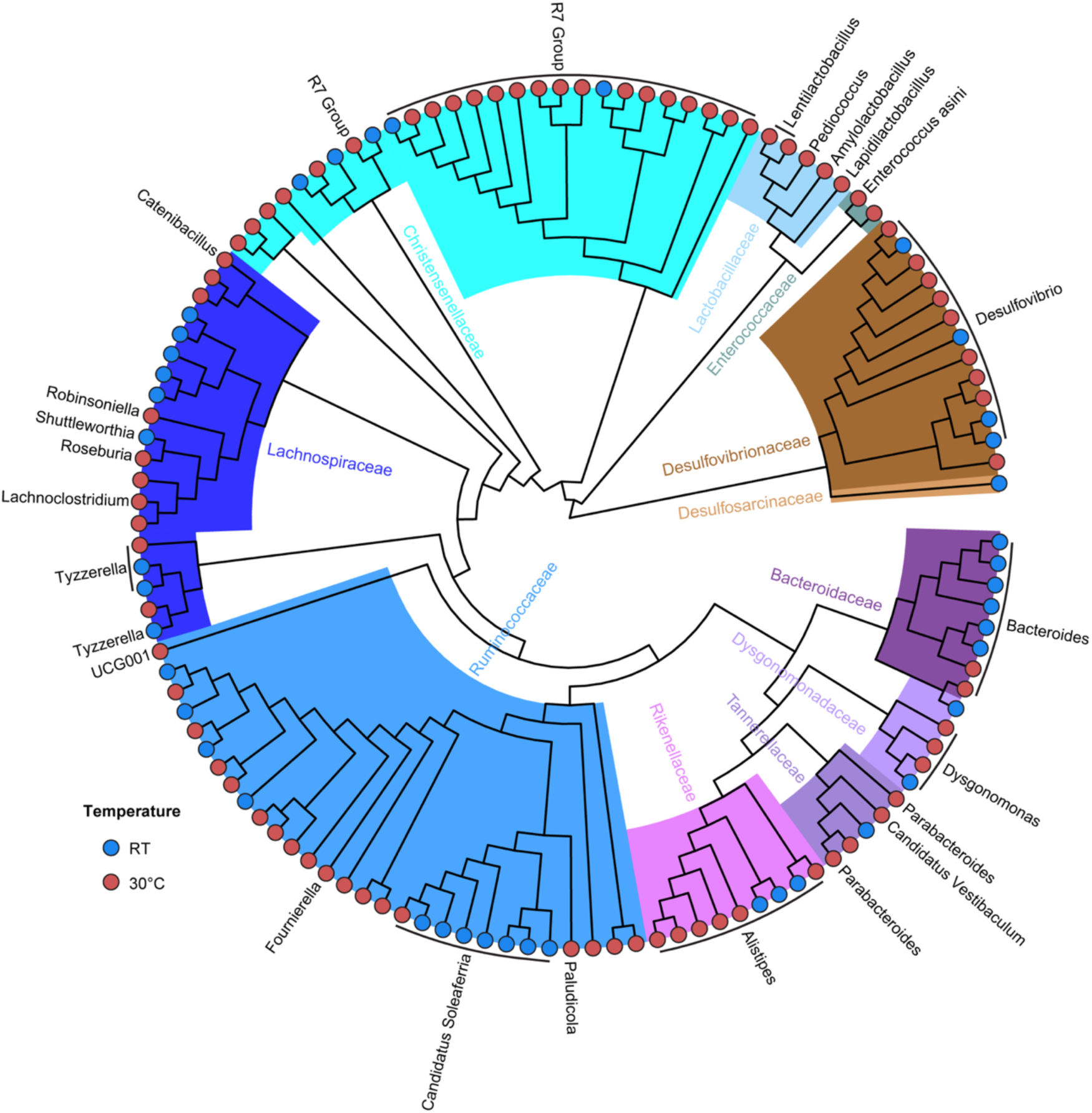
Phylogenetic tree of differentially abundant ASVs across temperature treatments. FastTree depicting differentially abundant ASVs in the most abundant families (maximum relative abundance > 10%). Highlights represent family level classifications. Leaf tip labels represent genus level classifications. ASVs without a leaf tip label were not classifiable at the genus level. Leaf tip colors depict which treatment ASVs were enriched in. Branch lengths not shown. RT = room temperature.

These ASV level differences translated into significant differences in family and phylum level abundances. At the family level, Wilcoxon rank sum tests found that Bacteroidaceae were significantly more abundant at room temperature (median difference = 2.07%, *p* < 0.001) while Lactobacillaceae and Enterococcaceae were significantly more abundant at 30°C (median difference = 1.94%, *p* = 0.01 and median difference = 0.41%, *p* = 0.005) (Figs 5A and S4). When summed at the phylum level, Bacteroidota and Firmicute (Bacillota) abundances were not significantly different across temperature treatments (*p* = 0.83 and *p* = 0.89). In contrast, the sulfate-reducing Desulfobacterota (Thermodesulfobacteridota) were significantly more abundant at room temperature (median difference = 3.5%, *p* = 0.04) while the Euryarchaeota (containing exclusively *Methanibrevibacter* ASVs) were significantly more abundant at 30°C (median difference = 0.17%, *p* = 0.005). Proteobacteria were significantly more abundant in the 30°C treatment (median difference = 0.26%, *p* = 0.04) (Figs 5B and S5).

**Fig 5.**
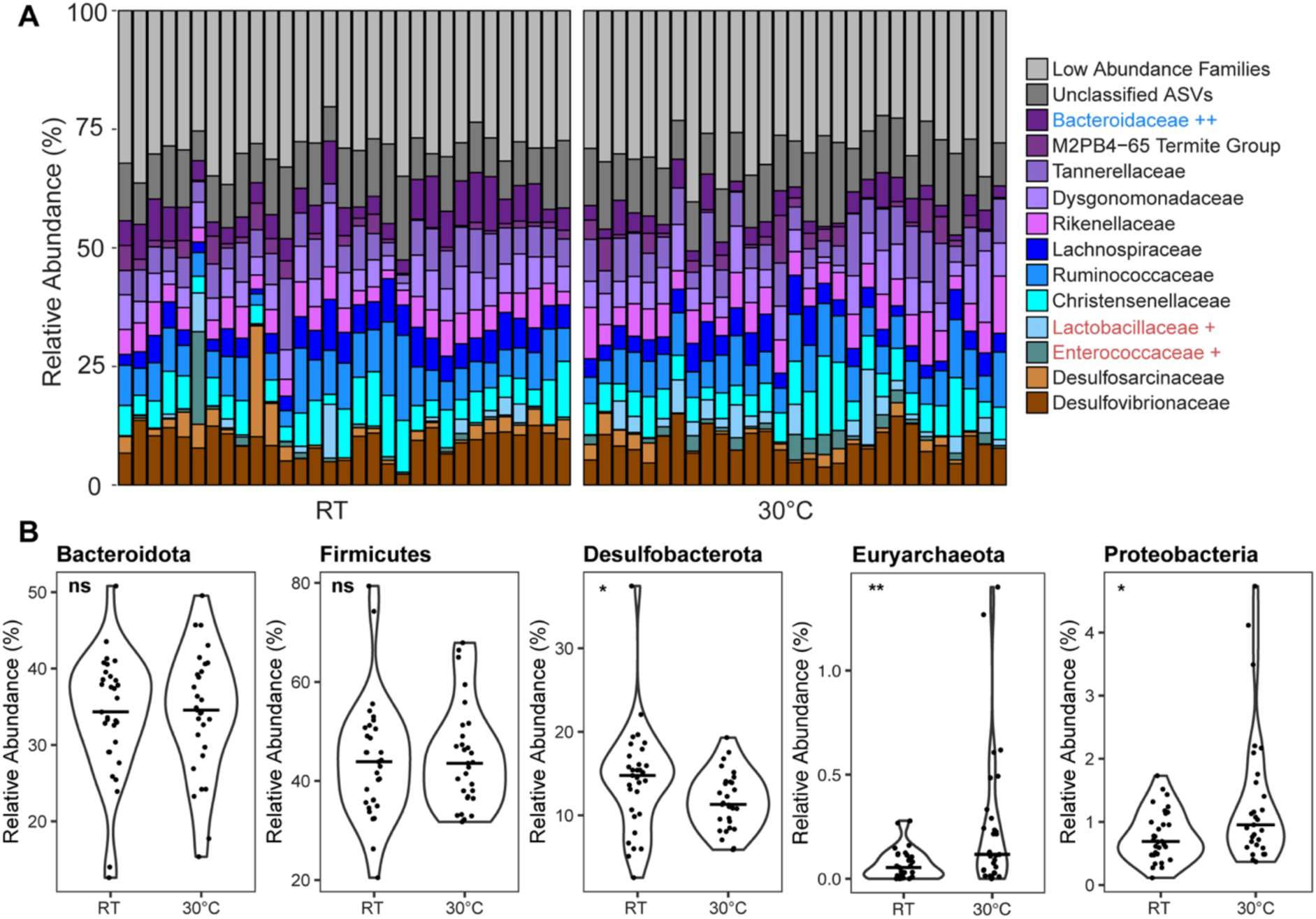
Comparison of family and phylum level relative abundances across temperature treatments. (A) Relative abundance of the 12 most abundant families (maximum relative abundance > 10%). Each bar represents an individual sample. ++ = higher in room temperature treatment, + = higher in 30°C treatment. (B) Violin plots showing the relative abundance of select phyla where bars represent the median and points represent individual samples. All libraries were resampled to a depth of 10,797 reads. Wilcoxon rank-sum tests were used to compare relative abundance across temperature treatments. RT = room temperature, * = *p* < 0.05, ** = *p* < 0.01, ns = no significance.

## Discussion

The goal of this study was to examine how changes in environmental temperature affect gut microbial community composition and stability in omnivorous cockroaches. We observed modest differences between room temperature (20-22°C) and 30°C treatments at the phylum and family level (Figs 5, S4, and S5). At the phylum level, there were significantly more Proteobacteria in the 30°C treatment than the room temperature treatment. The observed shift in Proteobacteria relative abundance was subtle, with a median difference of 0.26% (Fig 5). This response parallels studies in other insects and invertebrates. In a study of rough wood lice (*Porcellio scaber*), those maintained at 22°C had higher abundances of Proteobacteria (Enterobacteriaeae) compared to those maintained at 15°C (77). Experimental warming of the invertebrate nematode *Caenorhabditis elegans* also resulted in higher abundances of Proteobacteria (*Agrobacterium*) as temperature increased (15, 20, and 25°C treatments) (78). However, those studies observed more pronounced changes in Proteobacteria abundances compared to our findings in the American cockroach and were concurrent with substantial decreases in other common gut microbiota such as the Actinobacteria in *P. scaber* and *Sphingobacterium* in *C. elegans*. In the closely related termite *Reticulitermes flavipes*, Arango et al. saw a considerable increase in the abundance of Proteobacteria (*Acinetobacter,* Betaproteobacteriales, *Enterobacter*) at their lowest temperature treatment of 15°C compared to 27 or 35°C as well as subsequent decreases in Bacteroidota abundances (51). Studies in the fruit fly *Drosophila melanogaster* and cricket *Gryllus veletus* also report that Proteobacteria are responsive to temperature (79,80). However, the responsive Proteobacteria in the fruit fly and cricket studies are primarily *Wolbachia*, an intracellular bacterial symbiont that is not directly associated with the gut microbiome.

In vertebrate studies, increases in Proteobacteria abundance are associated with gut dysbiosis, pronounced shifts in community composition and diseases like irritable bowel syndrome and Crohn’s disease (81–83). The Proteobacteria are generally composed of opportunists that can survive a broad range of stressors compared to other phyla. It may be that changes in their abundance is indirectly determined by other community members response to temperature. In this scenario, Proteobacteria are well suited to ‘take over’ under conditions where more typical symbiotic populations are under environmental stress. We observed similar patterns with the Enterococcaceae and Lactobacillaceae in the 30°C treatment (Figs 4, 5 and S4), both of which are also commonly associated with gut dysbiosis (84,85). These taxa appear to bloom in a subset of our 30°C samples, consistent with our observation of increased beta-dispersion under high temperature conditions. This aligns with the “Anna Karenina” model where dysbiotic microbiota are unique and highly variable compared to healthy microbiota (86,87). We note a similar pattern in *Cephalotes rohweri* ant colonies exposed to high temperatures in field warming studies (27).

We also observed a significant decrease in Desulfobacterota at higher temperatures (Fig 5). The Desulfobacterota are highly active in *P. americana*, primarily acting as sulfate reducers (34,88). As sulfate reducers, they are likely in competition with methanogens for hydrogen (89,90). Cockroach gut methanogenic taxa include the Euryarchaeota which were significantly more abundant in the 30°C treatment (Fig 5). This suggests that hydrogen utilization may tip in favor of methanogens at higher temperatures, or that methanogens themselves fare better at higher temperatures, increasing hydrogen utilization and decreasing Desulfobacterota abundances. These observations align with studies in soils and batch reactors, where methanogens outcompete sulfate reducers as temperatures rise (91–93). Further studies are needed to better quantify the methanogenic archaea, as universal 16S rRNA primers used in this study are not optimized for their detection.

The modest shifts in gut microbiome composition that we report here contrasts with the experimental warming of *R. flavipes* termites, in which a shift from 27°C to 35°C resulted in large changes in phylum-level composition. Arango et al. reported that Bacteroidetes (Bacteroidota) almost doubled in relative abundance while Spirochaetes, Euryarchaeota and Elusimicrobia decreased. However, the authors noted that many of the microbial taxa lost following warming tended to be associated with ciliated protists that do not fare well at these higher temperatures (51). These protists are critical for breakdown of lignocellulosic material in lower termites (94). While similar protists have been reported in the cockroach, they are not essential for cockroach survival and energy acquisition (95). Microscopic observation of samples from our stock tanks indicate that a few protists are present but that they tend to be present at low abundance and low frequency. It may be that the ‘high’ temperature tested here was not high enough to negatively impact the protists, or that any reduction in protists had a negligible impact on overall community composition.

Together, these results suggest that the cockroach gut microbiome is relatively stable across a wide range of environmental temperatures. Although this resilience appears to be unique to cockroaches compared to other insect studies to date, comparable trends have been reported in wild *Anolis* lizards, where the gut microbiome was far less responsive to warming environmental temperature than in other lizards (96). While the 30°C temperature used in our study is within the preferred temperature range of this species, we observed that all cockroaches maintained at this temperature had a darker cuticle by the end of the experiment. Cuticle melanization has been directly linked to dessication tolerance in *Blattella germanica* (97), suggesting that our warming treatment was high enough to elicit a melanization stress response in *Periplaneta americana*. Additionally, it is common for bacteria to have lower tolerances for higher-than-preferred than lower-than-preferred temperatures (98). The high temperature tested shows some signs of increased variability and growth of opportunistic Proteobacteria, Enterococcaceae, and Lactobacillaceae which may have implications for the ability of these insects to serve as disease vectors in less climate-controlled settings. Finally, we highlight the potential for temperature to impact microbial competition between essential hydrogenotrophic organisms in the gut. This study suggests that the microbiome of ectotherms can be surprisingly resilient to temperature shifts of up to 10°C and has important implications for the ability of these insects to tolerate the impacts of climate change.

## Acknowledgements

This work was supported by the National Institute of General Medical Sciences of the National Institutes of Health under award number R35GM133789 and the National Science Foundation Graduate Research Fellowship Program under Grant No. 2236869. Any opinions, findings, and conclusions or recommendations expressed in this material are those of the author(s) and do not necessarily reflect the views of the National Science Foundation.

## Notes

### Competing Interest Statement

The authors have declared no competing interest.

### Summary of Updates

Figure references in the text were corrected, and minor changes were made to the discussion.

