## Supplemental figures S1 to S5 for "Large temperature excursions have modest impacts on community composition in the high diversity gut microbiome of omnivorous American cockroaches (*Periplaneta americana*)"

### Contents

**Description of supplementary tables** (uploaded as separate files) 2

**Fig S1:** Family level comparison of hindgut microbial community composition across temperature treatments 3

**Fig S2:** Comparison of between cohort Bray-Curtis dissimilarities 4

**Fig S3:** Heatmap of differentially abundant ASVs across temperature treatments 5

**Fig S4:** Family level relative abundances across temperature treatments 6

**Fig S5:** Phylum level relative abundances across temperature treatments 7

### Description of supplementary tables

**Table S1:** Bacterial load qPCR copy data and DNA extraction data.

**Table S2:** DESeq2 results table. Sheet one contains all results. The remaining sheets are split by the most abundant families (maximum relative abundance greater than 10%) and ASVs are filtered for significance ($p < 0.05$).

**Table S3:** Sample metadata and 16S rRNA amplicon read tracking data.

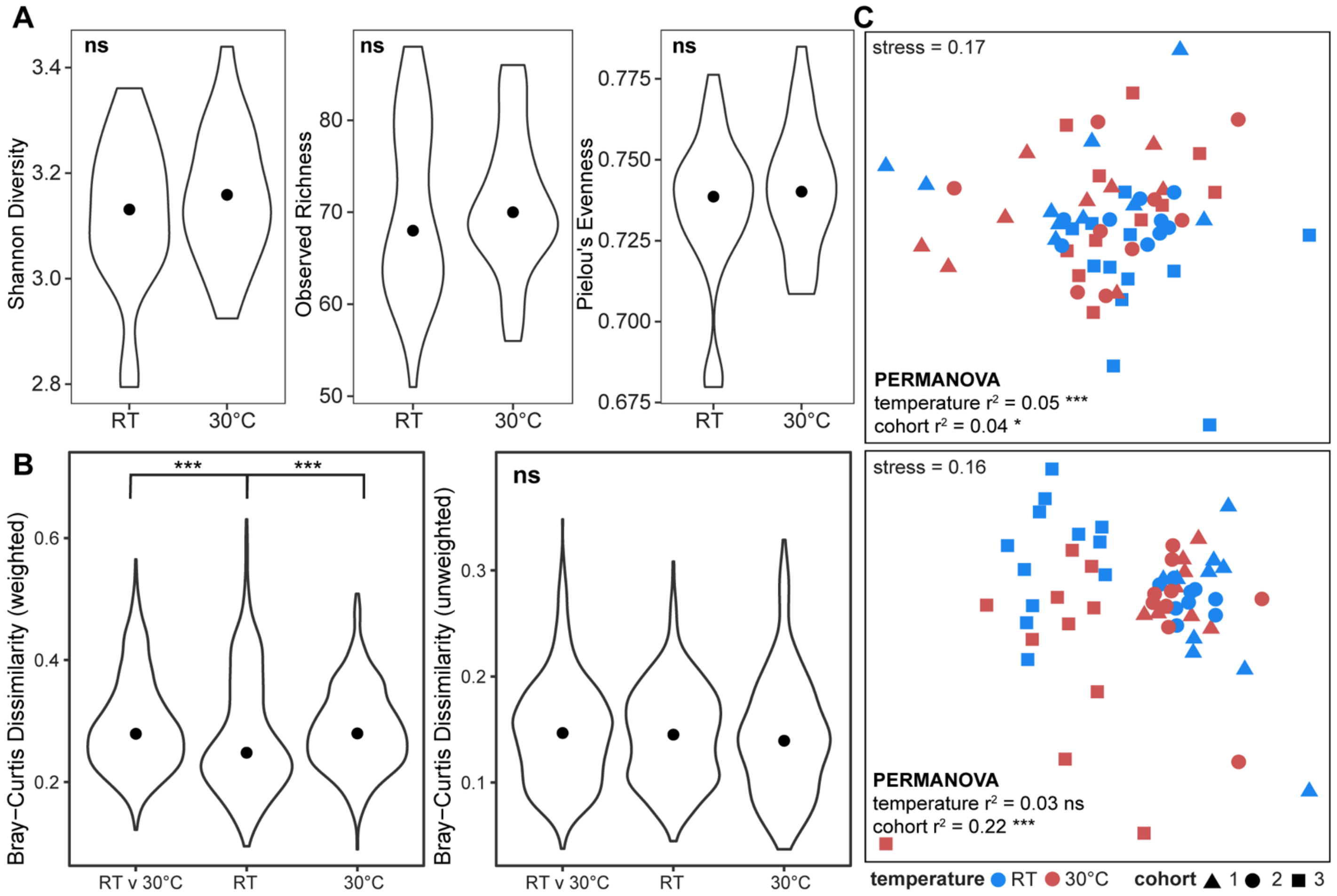


**Fig S1. Family level comparison of hindgut microbial community composition across temperature treatments.**

(A) Violin plots of alpha diversity measurements (Shannon diversity, observed richness, Pielou's evenness) where points represent the medians. (B) Violin plots of weighted (left) and unweighted (right) Bray-Curtis dissimilarities within and between temperature treatments where points represent the medians. Wilcoxon rank-sum tests were used to compare alpha diversity measures. Kruskal-Wallis and post-hoc Dunn's test with Bonferroni adjustment was used to compare Bray-Curtis dissimilarities. (C) Nonmetric multidimensional scaling (NMDS) of weighted (top) and unweighted (bottom) Bray-Curtis dissimilarities. NMDS stress was calculated with the metaMDS() function from the Vegan package. PERMANOVA was used to calculate $r^2$ and $p$ values for temperature and cohort. All libraries were resampled to a depth of 10,797 reads. RT = room temperature, * = $p < 0.05$, ** = $p < 0.01$, *** = $p < 0.001$, ns = no significance.

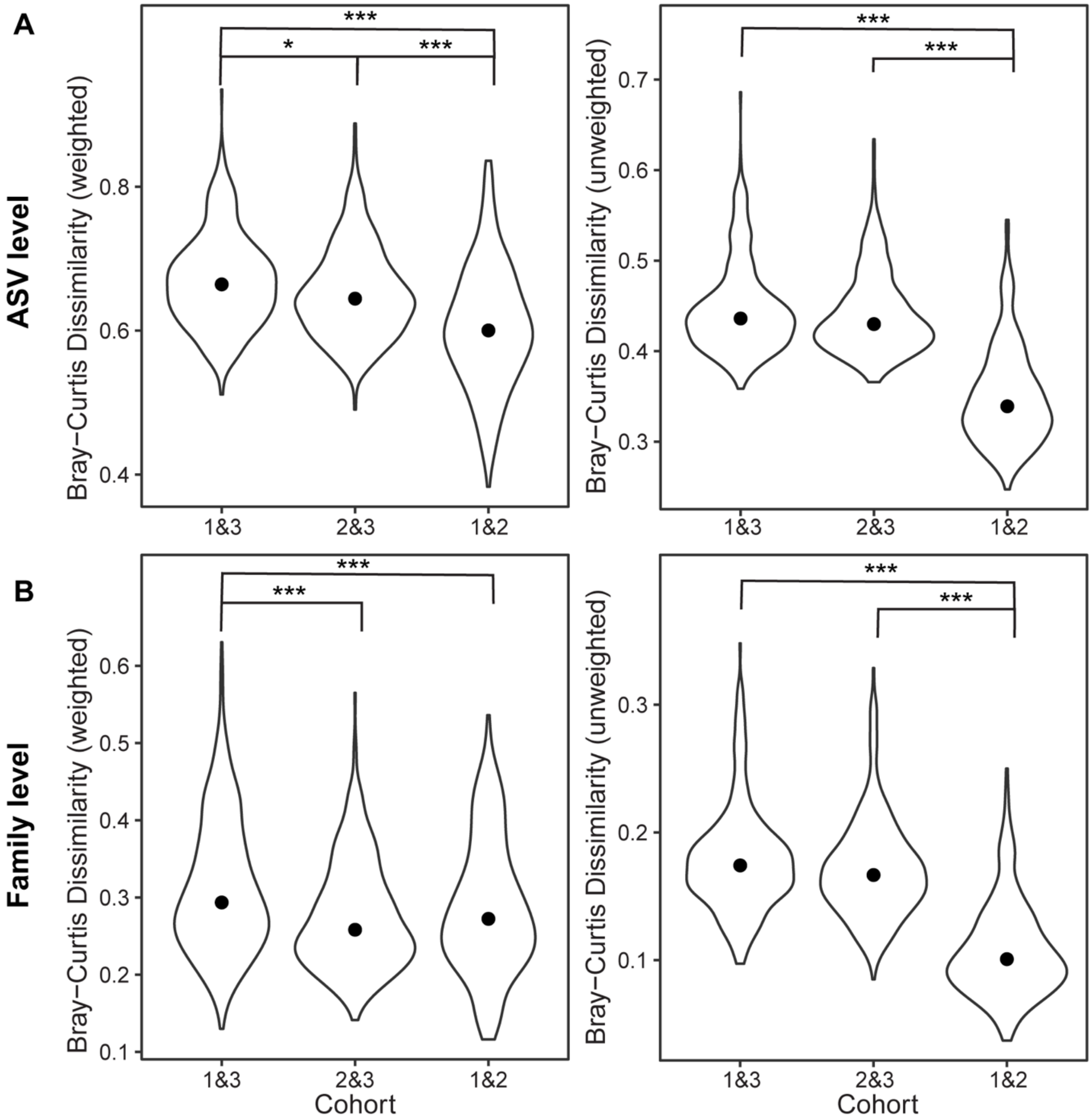


**Fig S2. Comparison of between cohort Bray-Curtis dissimilarities.**

Violin plots of weighted (left) and unweighted (right) Bray-Curtis dissimilarities between cohorts at the (A) ASV and (B) family level where points represent the medians. Kruskal-Wallis and post-hoc Dunn's test with Bonferroni adjustment was used to compare groups. All libraries were resampled to a depth of 10,797 reads. * = $p < 0.05$, *** = $p < 0.001$.

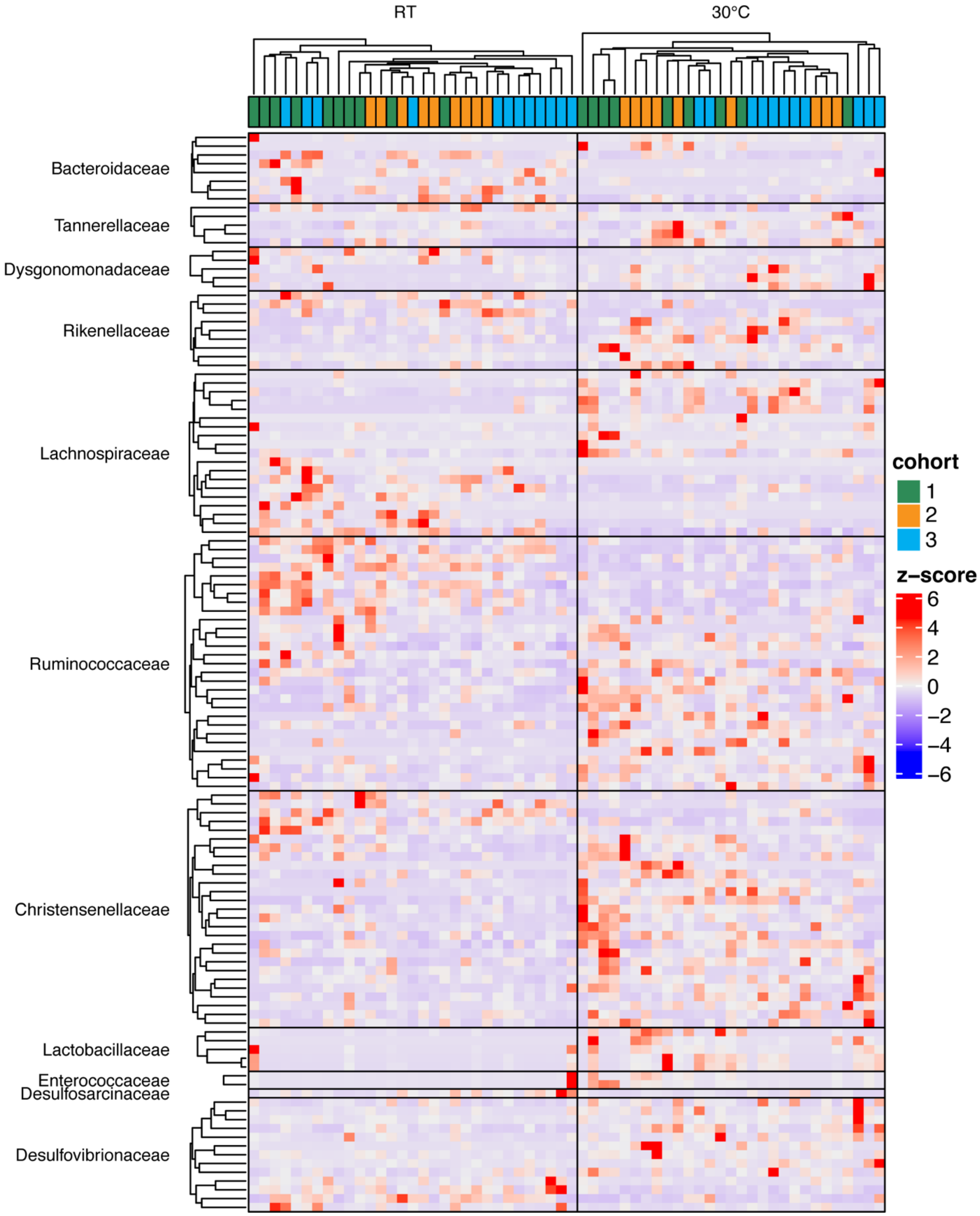


**Fig S3: Heatmap of differentially abundant ASVs across temperature treatments.**

Heatmap depicting the relative abundance of differentially abundant ASVs in the most abundant families (maximum relative abundance > 10%) as determined by DESeq2 (Table S2) ($p < 0.05$). Heatmap was generated using the ComplexHeatmap package with default clustering parameters. Relative abundances are Z-score standardized by row. RT = room temperature.

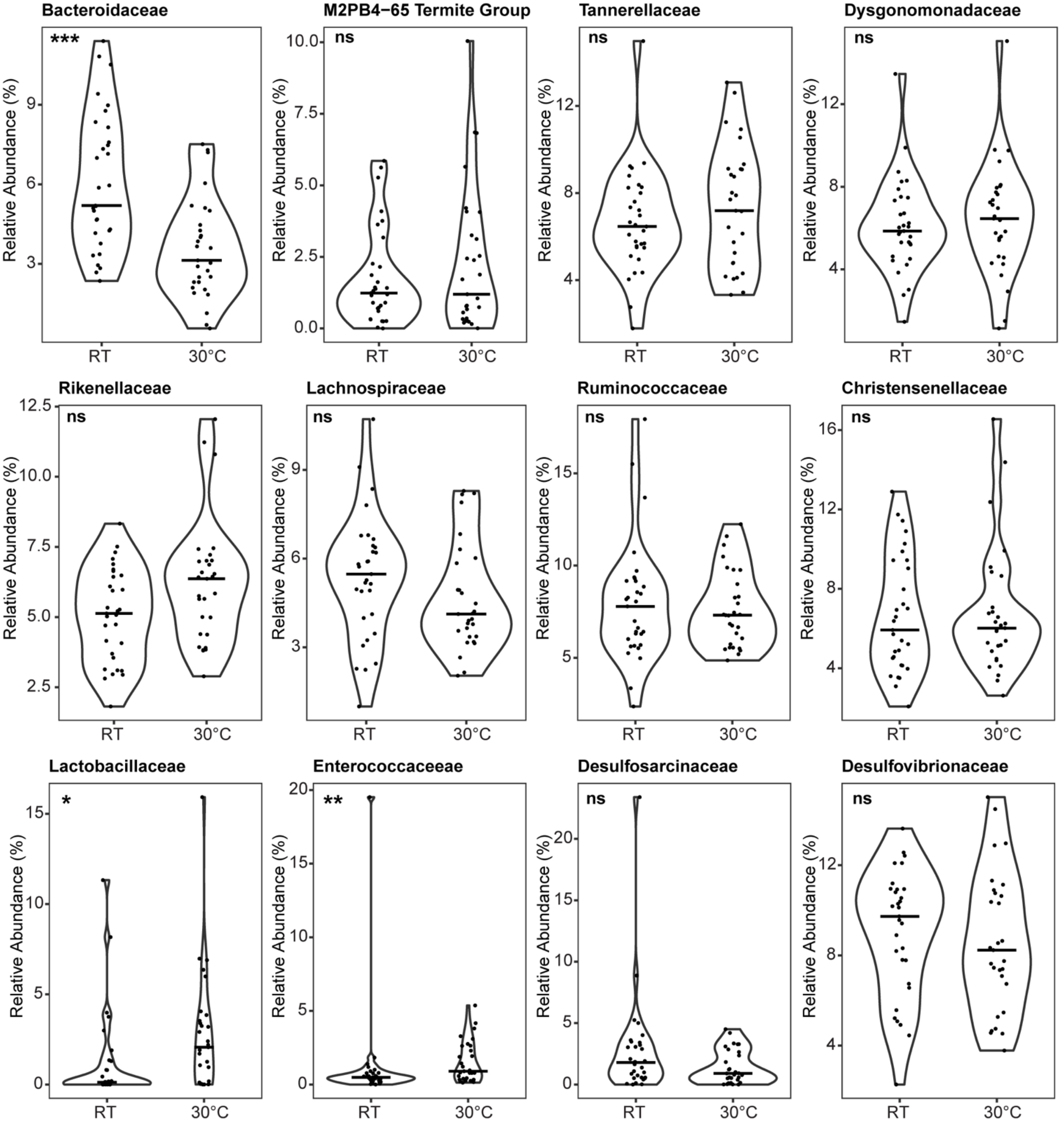


**Fig S4. Family level relative abundances across temperature treatments.**

Violin plots showing the relative abundances of the 12 most abundant families (maximum relative abundance > 10%). Bars represent the median and points represent individual samples. All libraries were resampled to a depth of 10,797 reads. Wilcoxon rank-sum tests were used to compare relative abundance across temperature treatments. RT = room temperature, * = $p < 0.05$, ** = $p < 0.01$, *** = $p < 0.001$, ns = no significance.

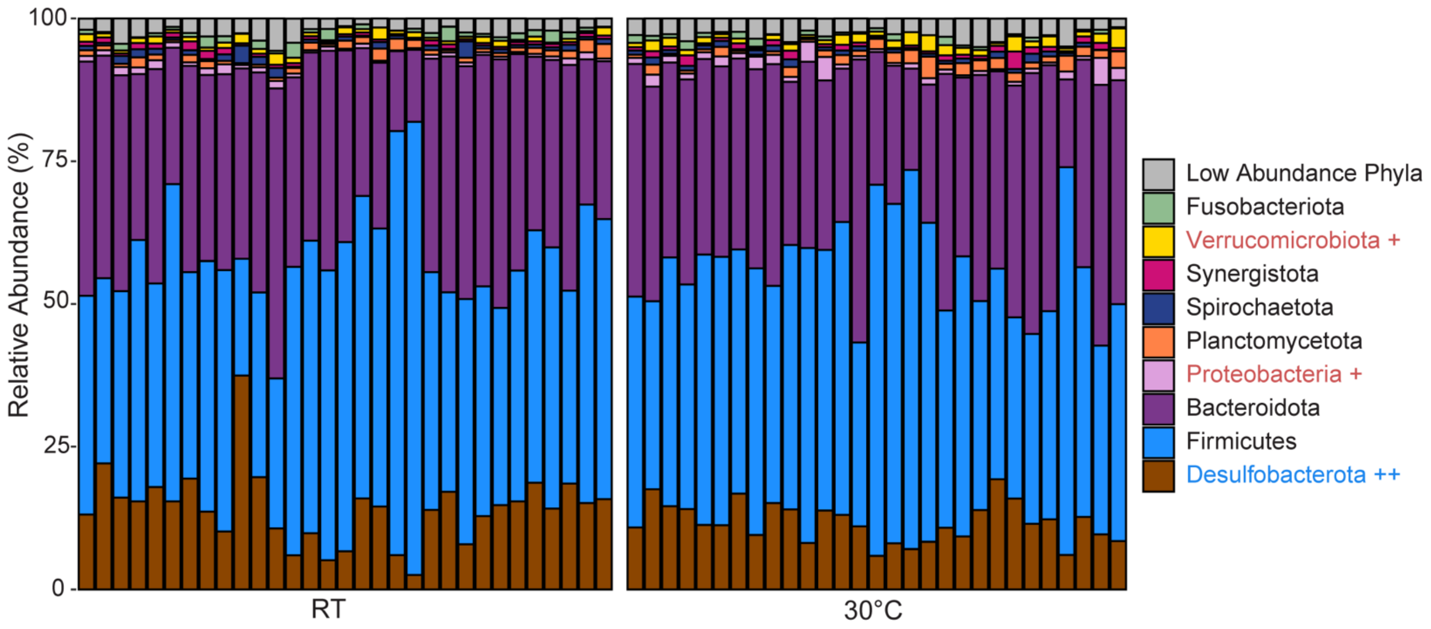


**Fig S5. Phylum level relative abundances across temperature treatments.**

Relative abundances of the 9 most abundant phyla (maximum relative abundance > 2%). Each stacked bar represents an individual sample. ++ = higher in room temperature treatment, + = higher in 30°C treatment as determined by Wilcoxon rank-sum tests ($p < 0.05$). RT = room temperature.
